# Quantitative Full-length transcriptome analysis by nanopore sequencing with Error-Aware UMI mapping

**DOI:** 10.64898/2026.01.14.699429

**Authors:** Qingwen Li, Dongxu Li, Chen Sun, Guangtao Song, Yihua Huang, Jizhong Lou

## Abstract

Comprehensive transcriptome profiling is essential for understanding RNA diversity and regulation, yet accurate identification and quantification of full-length transcript isoforms remain challenging with short-read sequencing technologies. Nanopore sequencing enables direct sequencing of long cDNA molecules and thus offers a powerful solution for full-length transcriptome analysis, but its application to quantitative transcriptomics is limited by PCR amplification bias and the difficulty of unique molecular identifier (UMI) recognition under high sequencing error rates.

Here, we developed UMImap, a dedicated pipeline for robust UMI identification, error correction, and deduplication in nanopore data. By integrating transcript-aware UMI correction with long-read isoform assembly, UMImap substantially improves UMI recognition accuracy compared with existing methods and effectively mitigates PCR-induced duplication bias. Using this framework, we identified tens of thousands of full-length transcript isoforms, including a large fraction of previously unannotated isoforms that are significantly longer than reference annotations.

Quantitative analyses demonstrate the reliability of UMImap for transcript-level quantification. Functional and pathway enrichment analyses of highly expressed novel isoforms revealed coherent and biologically meaningful patterns, including strong enrichment in RNA processing, splicing, and translation pathways.

Our results establish UMImap as an effective solution for UMI-based quantification in nanopore full-length transcriptome sequencing and highlight the potential of long-read sequencing to simultaneously achieve accurate isoform discovery and expression analysis in complex transcriptomes.

## Introduction

Transcriptome sequencing is a central tool in modern molecular biology, providing, enabling systematic investigation of RNA composition, regulation, and function across diverse biological contexts.[1–7]. At its core, transcriptome analysis aims to achieve two complementary objectives, accurate identification of transcript isoforms and reliable quantification of their expression level.[8–13] Isoform identification elucidates RNA diversity, including alternative splicing variants[14] and gene fusions,[15] while expression quantification measures transcript abundance to uncover gene regulation and functional mechanisms.[16, 17] Short-read next-generation sequencing (NGS) technologies have been widely adopted for transcriptome profiling studies due to their high throughput and accuracy.[18] However, the intrinsic limitations of short read length makes reconstruction of full-length transcripts highly challenging. Isoform inference from fragmented short reads relies on computational assembly, which often produces incomplete or ambiguous transcript models, particularly for genes with complex splicing patterns.[19, 20] Droplet-based platforms, such as Drop-seq[21] and 10X Genomics[22], further restrict sequencing to transcript termini, enabling accurate gene expression quantification but precluding comprehensive analysis of splicing patterns or full-length isoforms[23]. Even full-length protocols based on short reads, such as Smart-seq3,[24] remain constrained in their ability to accurately reconstruct very long or highly similar isoforms.

Long-read sequencing technologies have fundamentally changed this landscape. By generating reads that span entire RNA molecules, long-read platforms enable direct observation of full-length transcript structures without the need of assembly [25–27], thereby enabling precise isoform identification[28–31]. Nanopore sequencing, in particular, offers flexible library preparation, theoretically unlimited read length, and the capacity to capture complete isoforms in a single read. These features make nanopore sequencing especially well suited for transcript isoform discovery, characterization of alternative splicing, and detection of complex transcript variants such as gene fusions.

Accordingly, numerous computational tools have been developed for isoform identification and annotation using long-read transcriptome sequencing data. [14, 32, 33] These approaches have substantially expanded our understanding of RNA diversity across cell types and conditions. In contrast, transcript-level expression quantification using long-read sequencing remains comparatively underdeveloped.

Most existing quantification methods are tailored for direct RNA sequencing or assume negligible PCR bias in cDNA sequencing[16, 34]. Direct RNA sequencing suffers from the relatively high error rates of nanopore read, compromising accuracy[35, 36]. While in cDNA-based nanopore sequencing, PCR amplification is typically required, introducing duplication bias that can substantially distort transcript abundance estimates if not properly corrected.

Unique molecular identifiers (UMIs) provide a principled solution to PCR-induced bias by labeling individual molecules prior to amplification, enabling accurate identification and removal of duplicate reads. [11, 37–39] UMI-based strategies are now standard in short-read transcriptomics, including both bulk and single-cell applications. However, applying UMIs to nanopore sequencing presents unique challenges. The relatively high sequencing error rate complicates UMI recognition, as even small numbers of base errors can cause identical UMIs to appear distinct. Moreover, unlike sample barcodes, which are selected from predefined libraries such as the 96-barcode system used by Oxford Nanopore Technologies (ONT)[40, 41], UMIs are fully random sequences, UMIs are fully random sequences and lack an external reference, making error correction substantially more difficult.

Existing approaches to UMI handling in nanopore transcriptomics are limited. Some strategies rely on parallel short-read sequencing to generate high-accuracy UMI references for correcting nanopore sequencing results[38], increasing experimental complexity and cost. Tools originally developed for short-read UMI processing [42, 43] are generally incompatible with long-read data and nanopore-specific error profiles. Although dedicated tools such as Flexiplex[44] attempt to address these challenges, their performance in accurate UMI recognition and deduplication remains suboptimal, limiting the reliability of downstream quantification.

To overcome these limitations, we developed UMImap, a dedicated tool for UMI recognition, error-correction and deduplication pipeline specially designed for nanopore full-length transcriptome sequencing. UMImap leverages transcript-level mapping information to distinguish true UMIs from sequencing artifacts and applies alignment-based correction strategies tailored to nanopore error characteristics. Using the GM12878 cell line as a reference system, we established an integrated workflow that combines high-confidence isoform identification with robust, UMI-aware expression quantification.

In this study, we first constructed a comprehensive set of full-length transcript isoforms from nanopore cDNA sequencing data. We then applied UMImap to accurately identify and correct UMIs, enabling reliable transcript-level quantification across libraries generated with different PCR cycles. Finally, we performed functional and pathway enrichment analyses of highly expressed novel isoforms to assess their potential biological relevance. Together, our results demonstrate that UMImap effectively addresses a key bottleneck in nanopore transcriptomics and enables simultaneous, accurate analysis of transcript structure and expression, paves the way for the broader adoption of long-read nanopore sequencing in quantitative transcriptomics studies.

## Materials and Methods

### Cell Line, RNA Extraction, and cDNA Library Preparation

The human lymphoblastoid cell line GM12878 was used in this study. GM12878 is derived from a healthy female donor and is widely used as a reference sample in genomic and transcriptomic research[45–47]. Although immortalized with Epstein-Barr virus (EBV), GM12878 is generally considered representative of a non-cancerous human sample and has been extensively characterized [48].

Total RNA was extracted from GM12878 cells following standard protocols. Polyadenylated RNA molecules were reverse-transcribed into cDNA using oligo(dT) primers to specifically target mature mRNA transcripts. During reverse transcription, a template-switching oligonucleotide (TSO) primer containing a UMI sequence was incorporated at the 3’ end of the cDNA. This design ensured that each original RNA molecule was labeled with a unique UMI prior to PCR amplification.

Following cDNA synthesis, libraries were amplified by PCR. To evaluate the robustness of UMI-based deduplication under different amplification conditions, two libraries were generated from the same RNA input using 16 and 20 PCR cycles, respectively. The resulting libraries were prepared for nanopore sequencing according to the manufacturer’s standard cDNA sequencing protocols.

### Nanopore Sequencing and Raw Read Processing

Nanopore sequencing was performed using the PolyseqOne platform of Beijing Polyseq Biotech Inc. Raw sequencing reads were subjected to initial quality control and preprocessing prior to downstream analysis. Reads were first examined for strand orientation, as cDNA sequencing generates both sense and antisense reads.

To determine read orientation, local pairwise alignment based on the Smith-Waterman algorithm was employed to evaluate the similarity between the first 100 bases of each read and the expected primer sequences corresponding to the sense (TSO-derived) and antisense (oligo(dT)-derived) adapters. The scoring function is defined as:

*Smith-Waterman(i,j)*

where *s*(*a_i_,b_j_*) is the match score (+1) or mismatch penalty (-2), and *d* is the gap penalty, set to-1 for gap opening and-0.5 for gap extension. Reads with alignment scores below 13 for both adapters were discarded. Reads with higher scores for the sense adapter were identified as sense strands, with trailing primer sequences removed. Reads with higher scores for the antisense adapter were identified as antisense strands, they were reverse-complemented, and all adapter sequences were trimmed prior to further analysis.

### Preliminary UMI Segmentation

The workflow of UMImap is illustrated in Figure 1.

**Figure 1.**
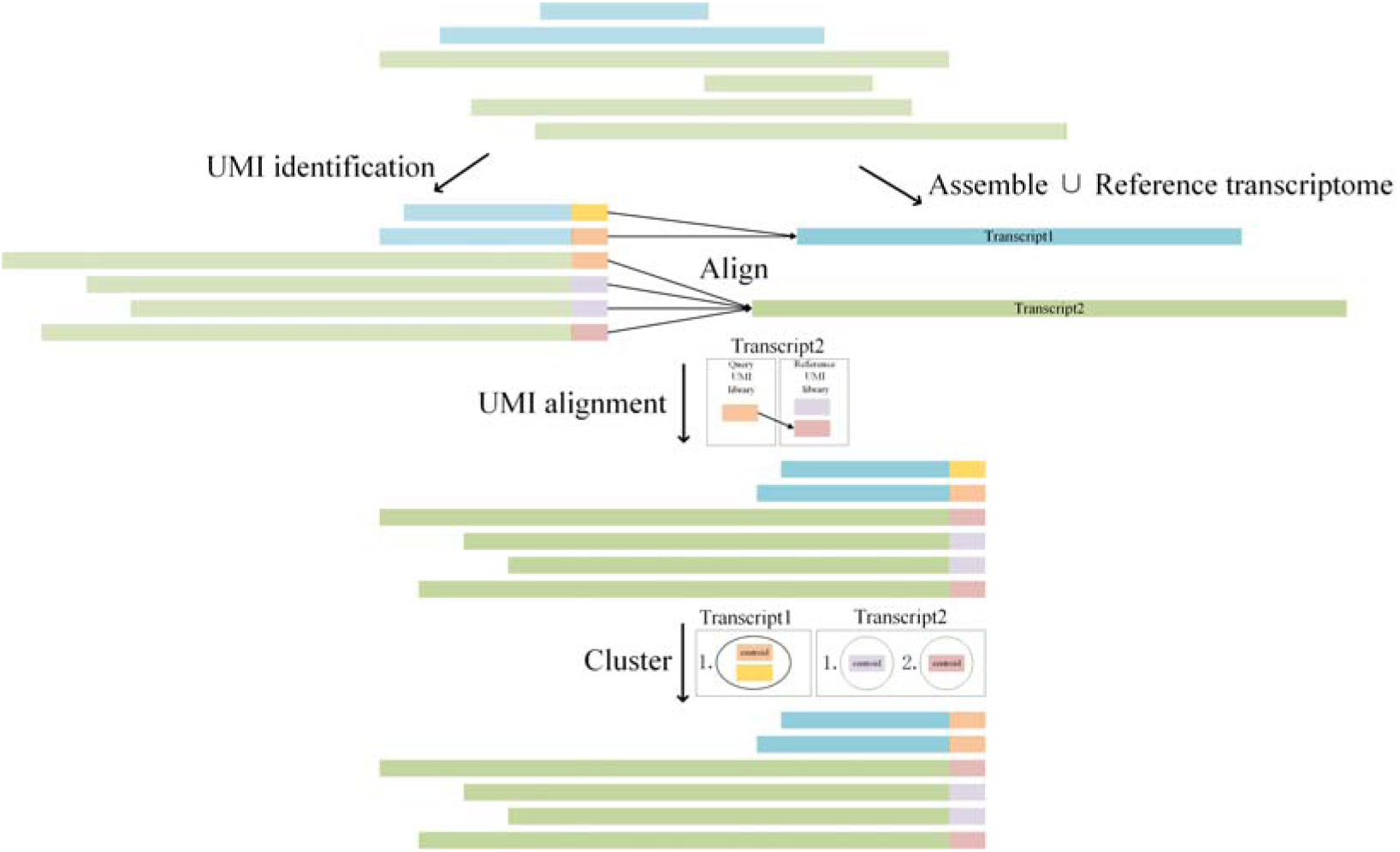
Overview of the UMImap pipeline for UMI-aware full-length transcriptome analysis. Schematic illustration of the UMImap workflow. Nanopore cDNA reads are first processed to determine strand orientation and trim adapter sequences. UMIs are preliminarily segmented based on flanking sequence patterns, followed by transcript isoform assembly and annotation. UMIs are then stratified into reference and query sets according to transcript mapping specificity. Transcript-aware alignment and error correction are applied to query UMIs, followed by UMI clustering and deduplication. The resulting corrected UMIs are used for accurate transcript-level expression quantification.

In this study, UMIs consisted of a 16-bp random nucleotide sequence flanked by fixed sequences derived from library design, including an 18-bp upstream sequence (CTTCCGATCTCAGCACCT) and an 8-bp downstream sequence (CTAATGGG). To locate UMI regions with each read, a sliding window approach was applied across the read sequence.

For each window, the Levenshtein distance between the read segment and the expected composite 42-bp sequence pattern (CTTCCGATCTCAGCACCTNNNNNNNNNNNNNNNNCTAATGGG)[44] was calculated. The Levenshtein distance is computed as:

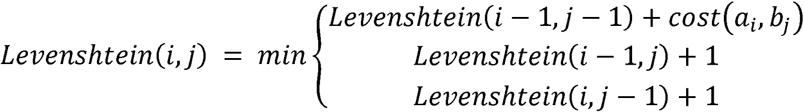

where *cost* (*a_i_,b_j_*)=0 if *a_i_ b_j_*, otherwise 1. The sliding window with the minimal Levenshtein distance was selected as the best match, and the central 16 bases were extracted as the preliminary UMI sequence. This step provided an initial UMI assignment while tolerating nanopore sequencing errors in the flanking regions.

### Transcript Isoform Assembly and Annotation

Full-length transcript isoforms were identified using the FLAIR pipeline[14]. Candidate isoforms were required to be supported by at least three reads, ensuring sufficient sequencing evidence for reliability. Additionally, supporting reads were required to cover at least 80% of the transcript length and to span a minimum of 25 bp into both the first and last exons to ensure structural completeness. The transcription start sites (TSS) and transcription end sites (TES) of supporting reads were required to cluster within a 100-bp window upstream and downstream, enforcing consistency in transcript boundaries.

The resulting transcript set were further evaluated using SQANTI3[49] for structural annotation and quality control. Transcripts fully matching known reference annotations were classified as full-splice matches (FSM) and were retained only if they showed no evidence of internal priming at the 3’ end, defined as an adenine proportion of <60% within the 20 bases downstream of the TES. Non-FSM transcripts were retained as high-confidence novel isoforms only if they lacked internal priming artifacts, showed no evidence of erroneous adapter sequences resulting from reverse transcription switching, and contained canonical splice junctions. These filters eliminated potential sequencing artifacts and identified novel isoforms absent from existing annotation databases. High-confidence novel isoforms were merged with reference annotations to generate a comprehensive transcriptome reference for downstream analyses.

### Construction of UMI Reference and Query Sets

In UMI-based transcriptome sequencing, reads originating from the same RNA molecule should map to a single transcript isoform. However, sequencing errors can cause misidentification of UMIs, leading to apparent mapping of the same UMI to multiple isoforms. To address this, UMIs were stratified based on their mapping behavior.

To optimize UMI identification, we constructed query and reference libraries for each transcript isoform. UMIs whose associated reads mapped exclusively to a single isoform were designated as high-confidence UMIs and used to construct a reference UMI set. UMIs whose reads mapped to multiple isoforms were placed into a query UMI set, representing candidates requiring error correction.

### UMI Alignment and Error Correction

UMIs in the query set were aligned to reference UMIs corresponding to the same transcript isoform using the Needleman-Wunsch global alignment algorithm[50], defined as:

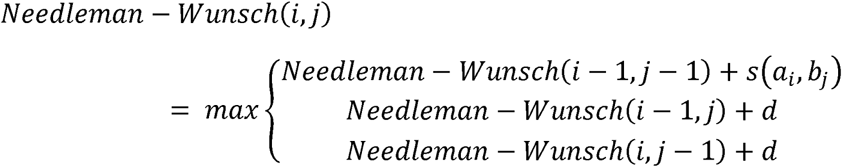

Where *s*(*a_i_,b_j_*) is a match score (+2) or a mismatch penalty (-4), and *d* is the gap penalty, set to-2 (deletion) or-20 (insertion) for gap opening and-1 (deletion) or-2 (insertion) for gap extension.

For each query UMI, the reference UMI with the highest similarity score was selected as a correction candidate. Query UMIs were corrected to the reference sequence if at least 12 matching positions were observed, thereby resolving sequence-induced errors while preserving true molecular diversity.

### UMI Clustering and Deduplication

UMI read depth was used as an indicator of library quality and UMI identification accuracy. A high proportion of UMIs supported by a single read is indicative of systematic identification errors. To reduce redundancy and further suppress error-derived UMIs, clustering analysis were performed on reference library UMIs for each transcript isoform [51]. Pairwise UMI similarity was calculated as the fraction of matching bases between UMI sequences:

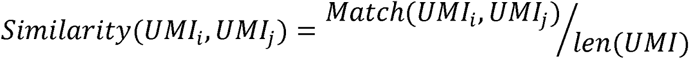

where *Math*(*UMI_i_,UMI_j_*) represents the number of matching bases between two UMIs. UMIs sharing greater than 80% sequence similarity were grouped into a single cluster, and the centroid sequence of each cluster was selected as the representative UMI sequence. All reads associated with UMIs in the same cluster were treated as originating from a single RNA molecule during downstream quantification.

### Expression Quantification and Downstream Analyses

Transcript expression levels were quantified by counting deduplicated UMIs assigned to each transcript isoform. Expression values were normalized to transcripts per million (TPM) to enable comparisons across samples.

For biological interpretation, novel isoforms with TPM ≥ 1 in both the 16-cycle and 20-cycle PCR libraries were selected for functional analysis. Gene Ontology (GO) enrichment and pathway enrichment analyses were performed using KEGG, Reactome, and WikiPathways databases. Statistical significance was assessed using standard enrichment frameworks, and results were interpreted in the context of known GM12878 biology.

## Results

### Evaluation of cDNA Library Quality

Full-length cDNA library quality is critical determinant of transcriptome analysis accuracy. Primers anneal to the 3’ poly(A) tail of RNA molecules during reverse transcription. However, RNA degradation and secondary structure can lead to premature termination of cDNA synthesis, resulting in reduced coverage toward TSS. Therefore, a gradual decrease in sequencing coverage from TES to TSS is an expected and widely accepted feature of high-quality full-length cDNA libraries.

To assess library quality, we calculated the normalized coverage per million reads (CPM) at each genomic position using deepTools2[52], examining read coverage across meta-genes as well as 1-kb regions flanking annotated TSS and TESBoth 16-cycle and 20-cycle PCR libraries exhibited highly consistent coverage patterns, with a smooth and progressive decline from TES to TSS (Figure 2a). The absence of abrupt coverage loss or positional bias indicates efficient reverse transcription and library construction. These results confirm that both libraries are suitable for downstream isoform discovery and quantitative analysis.

**Figure 2.**
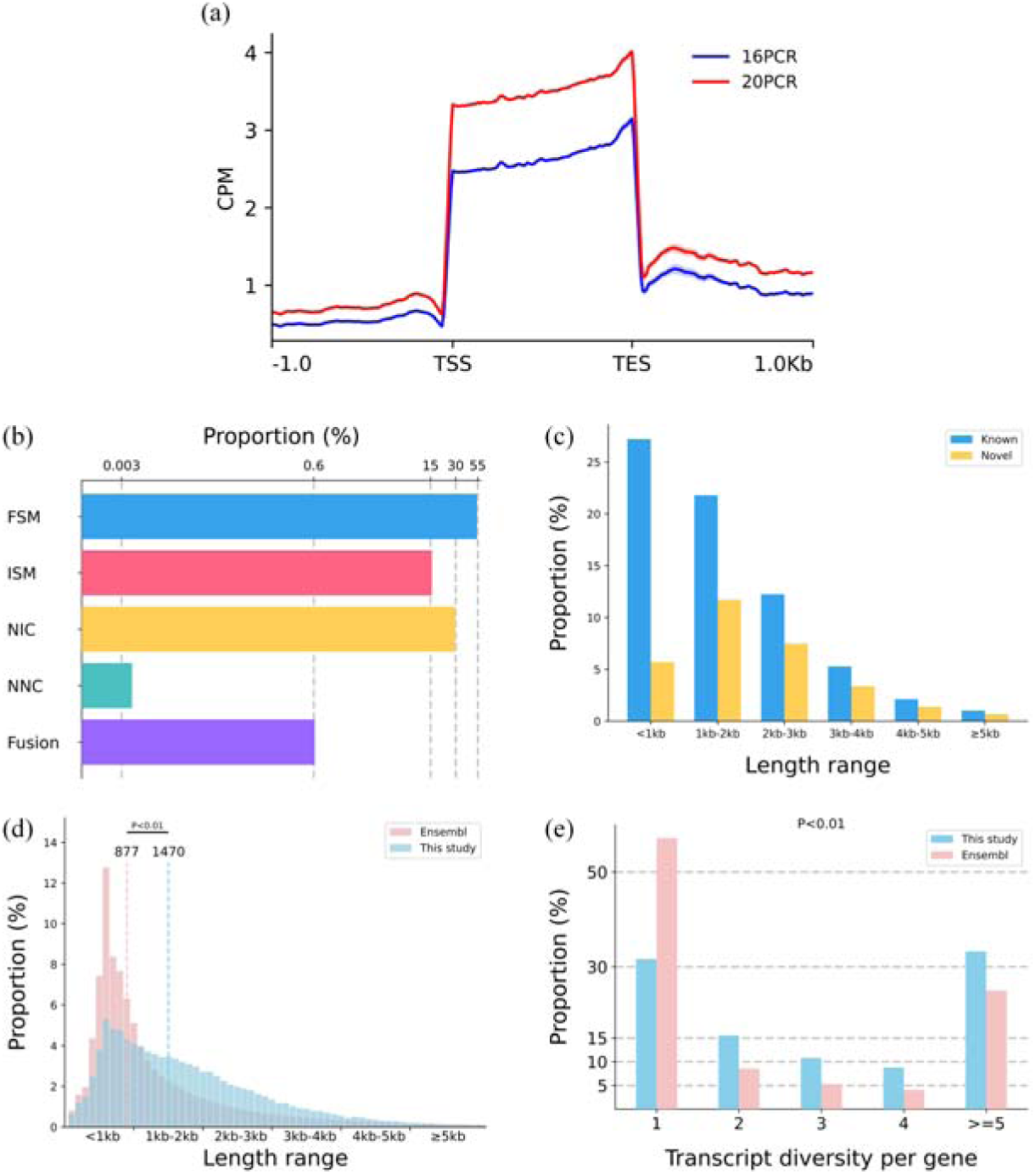
Quality assessment and structural characterization of nanopore-derived transcript isoforms. (a) Sequencing depth mapped to the genome for 16-cycle and 20-cycle PCR libraries. (b) Distribution of structural annotation categories for identified isoforms. (c) Length distribution of known and novel transcript isoforms. (d) Comparison of transcript length distributions between this study and Ensembl annotations. (e) Comparison of isoform diversity per gene between this study and Ensembl reference.

### Comprehensive Identification of Full-length Transcript Isoforms

A total of 75,030 transcript isoforms were identified, as supported by multiple reads and passing stringent structural quality filters. Structural annotation revealed that 69.68% of these isoforms corresponded transcripts present in the Ensembl reference database[53], including 54.08% classified as full-splice matches (FSM) and 15.60% as incomplete-splice matches (ISM) (Figure 2b). The remaining 30.32% represented previously unannotated isoforms, the majority of which (29.70%) were classified as novel in catalog (NIC), alongside a small fraction of fusion transcripts (0.62%), and novel not in catalog (NNC, 0.004%) isoforms.

Analysis of transcript length distribution highlighted clear differences between known and novel isoforms. Known isoforms were predominantly shorter than 1 kb, whereas novel isoforms were enriched in the 1–2 kb range (Figure 2c). When compared with Ensembl annotations, transcripts identified in this study exhibited a significantly longer median length ( 1,470 bp in this study versus 877 bp in Ensembl; independent t-test, P < 0.01) (Figure 2d). This shift toward longer transcript lengths underscores the advantage of nanopore long-read sequencing in capturing complete transcript structures that are often fragmented or missed by short-read approaches.

Beyond transcript length, nanopore sequencing substantially expanded isoform diversity at the gene level. Compared with Ensembl annotations, a significantly higher proportion of genes in our dataset harbored multiple isoforms, with more than 33.26% of genes exhibiting five or more distinct transcript variants (Chi-Squared test, P < 0.01) (Figure 2e). Together, these findings demonstrate that nanopore long-read sequencing enables comprehensive recovery of full-length transcript isoforms and reveals extensive RNA structural diversity, providing a powerful tool for understanding complex transcriptomic variability.

### Performance of UMImap in UMI Recognition

Accurate recognition of UMI is essential for correcting PCR amplification bias in nanopore cDNA sequencing. To evaluate the performance of UMImap, we compared it against UMI-tools[42], a widely used method developed for short-read sequencing, and Flexiplex[44], a tool specifically designed for analyzing of the nanopore sequencing data.

We first assessed transcript mapping specificity by calculating the proportion of reads associated with UMIs that mapped uniquely to a single transcript isoform. UMI-tools performed poorly in this context, with only 13.6% (16-cycle PCR) and 16.3% (20-cycle PCR) of reads mapping to a single transcript (Figure 3a), reflecting its incompatibility with nanopore error profiles. Flexiplex substantially improved mapping specificity, achieving 60.1% and 59.4% for the 16-cycle and 20-cycle PCR libraries respectively (Figure 3a), reflecting its ability in tailoring the systematic errors of nanopore sequencing. UMImap further increased these proportions to 67.3% and 66.2% (Figure 3a), demonstrating superior robustness in UMI recognition under high error-rate conditions.

**Figure 3.**
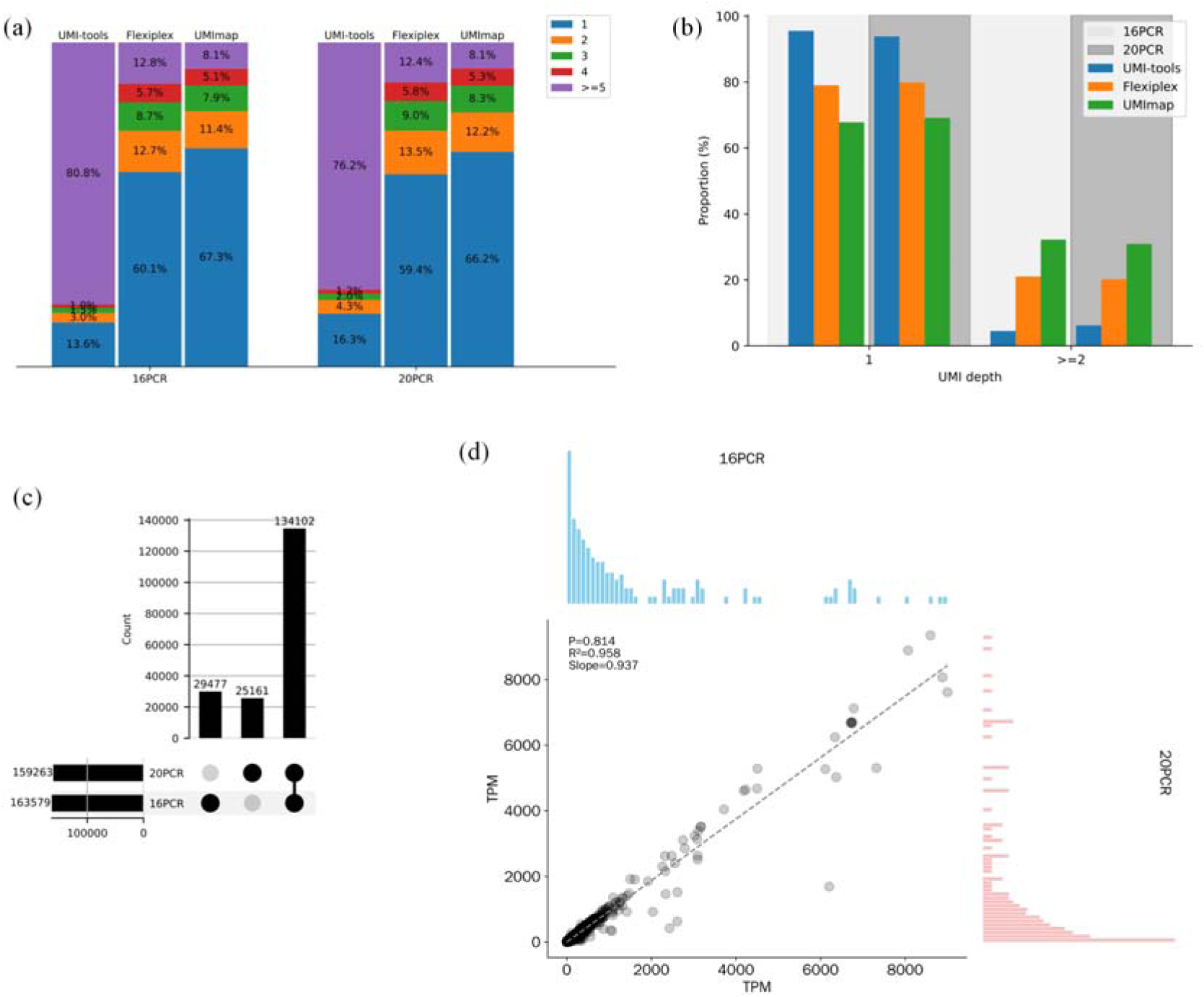
Performance of UMImap for UMI recognition and quantification. (a) Proportion of reads mapping uniquely to a single transcript using different UMI identification methods. (b) Comparison of the distribution of UMI read depths across different methods. (c) Number of expressed transcript isoforms detected in 16-cycle and 20-cycle PCR libraries. (d) Comparison of transcript expression levels between the 16-cycle and 20-cycle PCR libraries.

We next examined UMI read depth distributions, as erroneous UMI identification typically results in an excess of UMIs supported by only a single read.

UMImap consistently yielded the highest proportion of UMIs with read depths ≥2, reaching 32.2% in the 16-cycle PCR library and 30.9% in the 20-cycle PCR library (Figure 3b). These results indicate that UMImap more effectively consolidates reads originating from the same RNA molecule, reducing spurious UMI inflation and improving deduplication accuracy.

### Robust Transcript Quantification Across PCR Conditions

To assess whether UMImap effectively mitigates PCR amplification bias, we compared transcript expression estimates between libraries generated using different PCR cycle numbers (16-cycle and 20-cycle). Using UMImap-based deduplication, we detected expression of 163,579 transcript isoforms in the 16-cycle PCR library and 159,263 isoforms in the 20-cycle PCR library (Figure 3c). A total of 134,102 isoforms were shared between the two datasets and used for quantitative comparison. Transcript abundance was quantified as transcripts per million (TPM). The distributions of TPM values showed no significant differences between the two libraries (paired t-test, P = 0.814) (Figure 3d), despite the difference in PCR amplification depth. The high concordance demonstrates that UMImap effectively mitigates PCR-induced duplication biases and enables reproducible transcript-level quantification in nanopore full-length transcriptome sequencing.

### Functional and Pathway Enrichment of Novel Isoforms

To evaluate the biological relevance of newly identified transcripts, we focused on novel isoforms with TPM ≥ 1 in both libraries, yielding 15,553 isoforms corresponding to 5,689 genes. Functional annotation and enrichment analysis were performed to determine whether these isoforms represented coherent biological signals rather than technical artifacts.

Gene Ontology (GO) enrichment analysis[54] revealed highly consistent patterns across categories. Cellular components (CC) terms were strongly enriched for nuclear and nucleolar localization; molecular function (MF) terms were enriched for mRNA binding; and biological processes (BP) terms were dominated by RNA processing, mRNA splicing, and translation-related processes (Figure 4a). These results indicate that the identified novel isoforms are closely associated with core RNA regulatory machinery tightly coupled with important biological processes, confirming that these genes are functionally non-random.

**Figure 4.**
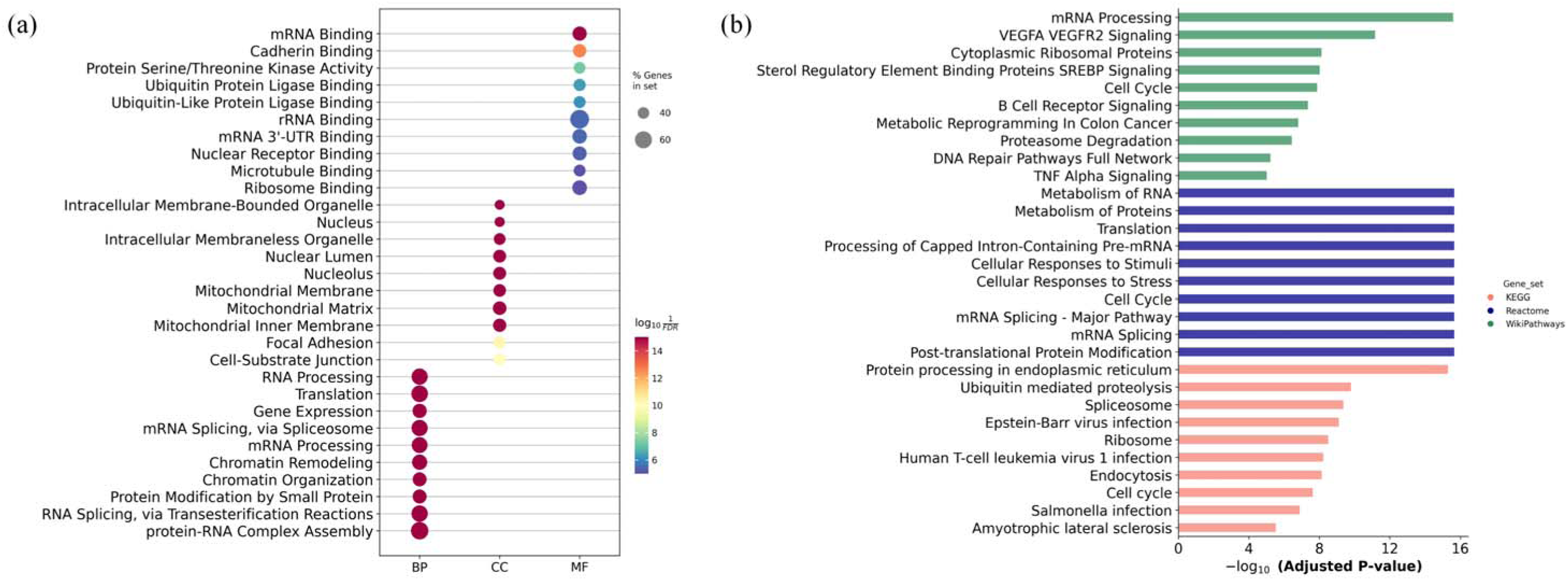
Functional enrichment of novel transcript isoforms. (a) GO enrichment analysis of genes corresponding to novel isoforms with TPM ≥ 1. (b) Pathway enrichment analysis using KEGG, Reactome, and WikePahtways databases.

Pathway enrichment analysis using KEGG[55], Reactome[56], and WikiPathways[57] databases further corroborated these findings. Across all these resources, the most significantly enriched pathways included the spliceosome, ribosome, and translation-related pathways (Figure 4b). The strong agreement among independent databases underscores the robustness of the enrichment results and minimizes the likelihood of database-specific bias.

The pathway enrichment results corroborated the GO findings, with RNA splicing and translation processes precisely mapped to molecular machinery such as the spliceosome and ribosome. This multi-level consistency, from individual gene functions to coordinated pathways, reinforces the biological significance of the newly identified transcript isoforms.

Notably, pathways related to Epstein-Barr virus infection were also significantly enriched, consistent with the EBV-immortalized background of GM12878 cell line. This concordance between enrichment results and known biological context further supports the authenticity of the identified novel isoforms.

Collectively, these functional analyses provide multi-level evidence – from gene function to pathway organization and biological context - that the novel isoforms uncovered by nanopore long-read sequencing and quantified using UMImap represent genuine components of the GM12878 transcriptome rather than random or technical artifacts.

## Discussions and Conclusions

Long-read sequencing technologies have transformed transcriptome analysis by enabling direct observation of full-length RNA molecules, thereby overcoming many limitations inherent to short-read sequencing. In this study, we demonstrate that nanopore cDNA sequencing, when combined with a dedicated and error-aware UMI processing strategy, can support not only comprehensive isoform discovery but also robust and reproducible transcript-level quantification. By developing UMImap and applying it to the GM12878 transcriptome, we address a critical bottleneck that has limited the broader adoption of nanopore sequencing in quantitative transcriptomics.

Our results highlight the intrinsic strengths of nanopore long-read sequencing for transcriptome analysis. Compared with reference annotations, the transcript isoforms identified in this study are substantially longer and exhibit markedly greater isoform diversity per gene. This expansion is not merely quantitative but qualitative, as long reads capture complete exon–intron structures and alternative splicing patterns that are difficult or impossible to reconstruct reliably from short-read data. The large proportion of novel isoforms identified, particularly those classified as novel in catalog, underscores the incompleteness of current transcript annotations and emphasizes the continued need for full-length transcriptome profiling in well-studied human cell lines.

Importantly, the structural features of the identified isoforms are supported by stringent filtering and quality control, as well as by coherent functional and pathway enrichment patterns. Together, these observations suggest that the novel isoforms detected here represent authentic components of the transcriptome rather than artifacts of sequencing or assembly. This finding reinforces the view that transcriptome complexity remains underappreciated, even in extensively characterized reference samples such as GM12878.

While nanopore sequencing excels at capturing transcript structure, accurate expression quantification has remained a major challenge. In cDNA-based nanopore sequencing, PCR amplification is unavoidable and introduces substantial duplication bias. Unique molecular identifiers offer a principled solution, but their effective use requires reliable UMI recognition and error correction in the presence of high sequencing error rates.

Our comparative analyses demonstrate that existing UMI processing tools are insufficient for nanopore transcriptome data. Methods originally designed for short-read sequencing fail to accommodate nanopore-specific error profiles, resulting in extensive UMI fragmentation and unreliable deduplication. Even tools explicitly developed for nanopore sequencing exhibit limitations in UMI recognition accuracy. These shortcomings directly translate into distorted transcript abundance estimates and undermine confidence in downstream analyses.

UMImap addresses this challenge by explicitly leveraging transcript-level mapping information and by applying alignment-based UMI correction strategies tailored to nanopore error characteristics. By distinguishing high-confidence UMIs from error-prone candidates and correcting the latter in a transcript-aware manner, UMImap substantially improves UMI consolidation and read deduplication. The resulting increase in UMI read depth and mapping specificity provides a direct measure of improved UMI recognition accuracy.

A key demonstration of UMImap’s effectiveness is the high concordance of transcript expression estimates between libraries generated with different PCR cycle numbers. Despite a substantial difference in amplification depth, expression profiles quantified using UMImap were statistically indistinguishable between the 16-cycle and 20-cycle libraries. This result provides strong empirical evidence that UMImap effectively mitigates PCR-induced duplication bias, enabling reproducible transcript-level quantification from nanopore cDNA sequencing data.

This robustness is particularly important for practical applications, where library preparation conditions may vary across experiments, batches, or laboratories. By reducing sensitivity to PCR amplification parameters, UMImap enhances the reliability and comparability of nanopore-based transcriptomic studies.

Beyond technical performance, the biological validity of the identified transcript isoforms is supported by multiple lines of evidence. Functional enrichment analyses consistently highlighted processes central to RNA biology, including RNA binding, splicing, and translation, reflecting the expected functional landscape of a transcriptionally active human cell line. The convergence of enrichment results across independent pathway databases further strengthens confidence in these findings.

Notably, enrichment of pathways related to Epstein–Barr virus infection aligns with the known biological background of GM12878 cells, which are EBV-immortalized. This concordance between sequencing-derived results and established biological context argues strongly against the notion that the observed novel isoforms arise from technical noise. Instead, it suggests that long-read sequencing coupled with accurate UMI-based quantification can sensitively capture condition-and cell-type–specific transcriptomic features.

Despite these advances, several limitations warrant consideration. First, while UMImap substantially improves UMI recognition, extremely high sequencing error rates or very short UMI designs may still pose challenges. Future improvements in nanopore basecalling accuracy and library design are likely to further enhance UMI-based quantification. Second, this study focused on bulk transcriptome sequencing of a single reference cell line. Extending UMImap to more diverse biological samples, including primary tissues and single-cell applications, will be an important direction for future work.

In addition, while nanopore sequencing offers unparalleled flexibility and read length, its throughput and cost structure differ from those of short-read platforms. Hybrid strategies that integrate nanopore long reads with short-read sequencing may further improve transcriptome characterization and quantification, particularly for large-scale studies.

In summary, this study establishes a robust framework for full-length transcriptome analysis using nanopore sequencing. By integrating high-confidence isoform identification with UMImap-based UMI recognition and deduplication, we demonstrate that nanopore sequencing can support accurate and reproducible transcript-level quantification, despite inherent sequencing errors and PCR amplification biases. The large number of biologically meaningful novel isoforms identified here underscores the power of long-read technologies to reveal previously hidden layers of transcriptome complexity. UMImap directly addresses a key limitation of nanopore transcriptomics and provides a practical solution for quantitative full-length RNA sequencing. As nanopore sequencing continues to mature, tools such as UMImap will be essential for unlocking its full potential in transcriptomics, enabling comprehensive and reliable analysis of RNA structure and expression across a wide range of biological and biomedical applications.

## Acknowledgements

This work is partly supported by grants from the Ministry of Science and Technology of China (2024YFC3406200) and the Strategic Priority Research Program of the Chinese Academy of Sciences (XDB1000103).

## Conflict of Interests

Guangtao Song, Yihua Huang and Jizhong Lou are co-founders and shareholders of Beijing Polyseq Biotech Co. Ltd. Beijing Polyseq Biotech Co. Ltd. and Institute of Biophysics, Chinese Academy of Sciences have filed a patent using materials described in this article.

## Software and data Availability

The code of UMImap can be accessed from Github (https://github.com/liqingwen98/UMImap).

## Notes

https://github.com/liqingwen98/UMImap

### Competing Interest Statement

The authors have declared no competing interest.

## References

1. Conesa, A., et al., A survey of best practices for RNA-seq data analysis. Genome Biol, 2016. 17: p. 13.

2. Han, X., et al., Construction of a human cell landscape at single-cell level. Nature, 2020. 581(7808): p. 303–309.

3. Li, Q., et al., Prediction of Anticancer Peptides Using a Low-Dimensional Feature Model. Frontiers in Bioengineering and Biotechnology, 2020. Volume 8 - 2020.

4. Li, Q., et al., Identification and classification of promoters using the attention mechanism based on long short-term memory. Frontiers of Computer Science, 2022. 16(4): p. 164348.

5. Li, Q., et al., Identification and Classification of Enhancers Using Dimension Reduction Technique and Recurrent Neural Network. Computational and Mathematical Methods in Medicine, 2020. 2020(1): p. 8852258.

6. Li, Q., et al., Identification of Secreted Proteins From Malaria Protozoa With Few Features. IEEE Access, 2020. 8: p. 89793–89801.

7. Qiao, J., et al., Towards Retraining-free RNA Modification Prediction with Incremental Learning. Information Sciences, 2024: p. 120105.

8. Loving, R., et al., Long-read sequencing transcriptome quantification with lr-kallisto. bioRxiv, 2024: p. 2024.07.19.604364.

9. Byrne, A., et al., Nanopore long-read RNAseq reveals widespread transcriptional variation among the surface receptors of individual B cells. Nat Commun, 2017. 8: p. 16027.

10. Volden, R. and C. Vollmers, Single-cell isoform analysis in human immune cells. Genome Biol, 2022. 23(1): p. 47.

11. Rebboah, E., et al., Mapping and modeling the genomic basis of differential RNA isoform expression at single-cell resolution with LR-Split-seq. Genome Biol, 2021. 22(1): p. 286.

12. Fan, X., et al., Single-cell RNA-seq analysis of mouse preimplantation embryos by third-generation sequencing. PLoS Biology, 2020. 18(12): p. e3001017.

13. Wei, L., et al., Improved and Promising Identification of Human MicroRNAs by Incorporating a High-Quality Negative Set. IEEE/ACM Transactions on Computational Biology and Bioinformatics, 2014. 11(1): p. 192–201.

14. Tang, A.D., et al., Full-length transcript characterization of SF3B1 mutation in chronic lymphocytic leukemia reveals downregulation of retained introns. Nature Communications, 2020. 11(1): p. 1438.

15. Chen, Y., et al., Gene Fusion Detection and Characterization in Long-Read Cancer Transcriptome Sequencing Data with FusionSeeker. Cancer Res, 2023. 83(1): p. 28–33.

16. Zare Jousheghani, Z., N.P. Singh, and R. Patro, Oarfish: enhanced probabilistic modeling leads to improved accuracy in long read transcriptome quantification. Bioinformatics, 2025. 41(Supplement_1): p. i304-i313.

17. Jin, J., et al., iDNA-ABF: multi-scale deep biological language learning model for the interpretable prediction of DNA methylations. Genome biology, 2022. 23(1): p. 1–23.

18. Tilgner, H., et al., Comprehensive transcriptome analysis using synthetic long-read sequencing reveals molecular co-association of distant splicing events. Nature Biotechnology, 2015. 33(7): p. 736–742.

19. Baldoni, P.L., et al., Dividing out quantification uncertainty allows efficient assessment of differential transcript expression with edgeR. Nucleic Acids Res, 2024. 52(3): p. e13.

20. Zhu, A., et al., Nonparametric expression analysis using inferential replicate counts. Nucleic Acids Res, 2019. 47(18): p. e105.

21. Zheng, G.X., et al., Massively parallel digital transcriptional profiling of single cells. Nat Commun, 2017. 8: p. 14049.

22. Coombe, L., et al., Assembly of the Complete Sitka Spruce Chloroplast Genome Using 10X Genomics’ GemCode Sequencing Data. PLoS One, 2016. 11(9): p. e0163059.

23. Arzalluz-Luque, Á. and A. Conesa, Single-cell RNAseq for the study of isoforms-how is that possible? Genome Biol, 2018. 19(1): p. 110.

24. Hagemann-Jensen, M., et al., Single-cell RNA counting at allele and isoform resolution using Smart-seq3. Nat Biotechnol, 2020. 38(6): p. 708–714.

25. De Paoli-Iseppi, R., J. Gleeson, and M.B. Clark, Isoform Age - Splice Isoform Profiling Using Long-Read Technologies. Front Mol Biosci, 2021. 8: p. 711733.

26. Li, Q., et al., BaseNet: A transformer-based toolkit for nanopore sequencing signal decoding. Comput Struct Biotechnol J, 2024. 23: p. 3430–3444.

27. Li, Q., et al., RMNet: An RNA m6A Cross-species Methylation Detection Method for Nanopore Sequencing. Curr Drug Targets, 2025.

28. Sharon, D., et al., A single-molecule long-read survey of the human transcriptome. Nat Biotechnol, 2013. 31(11): p. 1009–14.

29. Soneson, C., et al., A comprehensive examination of Nanopore native RNA sequencing for characterization of complex transcriptomes. Nat Commun, 2019. 10(1): p. 3359.

30. Weirather, J.L., et al., Comprehensive comparison of Pacific Biosciences and Oxford Nanopore Technologies and their applications to transcriptome analysis. F1000Res, 2017. 6: p. 100.

31. Yin, C., et al., NanoCon: contrastive learning-based deep hybrid network for nanopore methylation detection. Bioinformatics, 2024. 40(2): p. btae046.

32. Lienhard, M., et al., IsoTools: a flexible workflow for long-read transcriptome sequencing analysis. Bioinformatics, 2023. 39(6).

33. Wyman, D., et al., A technology-agnostic long-read analysis pipeline for transcriptome discovery and quantification. bioRxiv, 2020: p. 672931.

34. Gleeson, J., et al., Accurate expression quantification from nanopore direct RNA sequencing with NanoCount. Nucleic Acids Res, 2022. 50(4): p. e19.

35. Neumann, D., A.S. Reddy, and A. Ben-Hur, RODAN: a fully convolutional architecture for basecalling nanopore RNA sequencing data. BMC bioinformatics, 2022. 23(1): p. 142.

36. Li, Q., et al., GCRTcall: a transformer based basecaller for nanopore RNA sequencing enhanced by gated convolution and relative position embedding via joint loss training. Frontiers in Genetics, 2024. Volume 15 - 2024.

37. Ebrahimi, G., et al., Fast and accurate matching of cellular barcodes across short-reads and long-reads of single-cell RNA-seq experiments. iScience, 2022. 25(7): p. 104530.

38. Lebrigand, K., et al., High throughput error corrected Nanopore single cell transcriptome sequencing. Nat Commun, 2020. 11(1): p. 4025.

39. Long, Y., et al., FlsnRNA-seq: protoplasting-free full-length single-nucleus RNA profiling in plants. Genome Biol, 2021. 22(1): p. 66.

40. Dommann, J., et al., A novel barcoded nanopore sequencing workflow of high-quality, full-length bacterial 16S amplicons for taxonomic annotation of bacterial isolates and complex microbial communities. mSystems, 2024. 9(10): p. e00859–24.

41. Technologies, O.N. Native Barcoding Kit 96 V14. 2025; Available from: https://nanoporetech.com/document/ligation-sequencing-gdna-native-barcoding-v14-sqk-nbd114-96.

42. Smith, T., A. Heger, and I. Sudbery, UMI-tools: modeling sequencing errors in Unique Molecular Identifiers to improve quantification accuracy. Genome Res, 2017. 27(3): p. 491–499.

43. Tsagiopoulou, M., et al., UMIc: A Preprocessing Method for UMI Deduplication and Reads Correction. Frontiers in Genetics, 2021. Volume 12 - 2021.

44. Cheng, O., et al., Flexiplex: a versatile demultiplexer and search tool for omics data. Bioinformatics, 2024. 40(3).

45. Gibbs, R.A., et al., The International HapMap Project. Nature, 2003. 426(6968): p. 789-796.

46. Jain, M., et al., Nanopore sequencing and assembly of a human genome with ultra-long reads. Nat Biotechnol, 2018. 36(4): p. 338–345.

47. Wang, R., et al., DeepBIO: an automated and interpretable deep-learning platform for high-throughput biological sequence prediction, functional annotation and visualization analysis. Nucleic Acids Research, 2023. 51(7): p. 3017–3029.

48. Dunham, I., et al., An integrated encyclopedia of DNA elements in the human genome. Nature, 2012. 489(7414): p. 57-74.

49. Pardo-Palacios, F.J., et al., SQANTI3: curation of long-read transcriptomes for accurate identification of known and novel isoforms. Nature Methods, 2024. 21(5): p. 793–797.

50. Rognes, T., et al., VSEARCH: a versatile open source tool for metagenomics. PeerJ, 2016. 4: p. e2584.

51. Fu, L., et al., CD-HIT: accelerated for clustering the next-generation sequencing data. Bioinformatics, 2012. 28(23): p. 3150–2.

52. Ramírez, F., et al., deepTools2: a next generation web server for deep-sequencing data analysis. Nucleic Acids Research, 2016. 44(W1): p. W160–W165.

53. Dyer, S.C., et al., Ensembl 2025. Nucleic Acids Research, 2024. 53(D1): p. D948–D957.

54. Consortium, T.G.O., The Gene Ontology resource: enriching a GOld mine. Nucleic Acids Research, 2020. 49(D1): p. D325–D334.

55. Kanehisa, M., et al., KEGG: biological systems database as a model of the real world. Nucleic Acids Research, 2024. 53(D1): p. D672–D677.

56. Fabregat, A., et al., The Reactome Pathway Knowledgebase. Nucleic Acids Research, 2017. 46(D1): p. D649–D655.

57. Agrawal, A., et al., WikiPathways 2024: next generation pathway database. Nucleic Acids Research, 2023. 52(D1): p. D679–D689.

